# Effects of Prospective Motion Correction on Perivascular Spaces at 7T MRI Evaluated Using Motion Artifact Simulation

**DOI:** 10.1101/2023.12.25.573328

**Authors:** Bingbing Zhao, Yichen Zhou, Xiaopeng Zong

## Abstract

**Purpose:** Prospective motion correction (PMC) is a promising method in mitigating motion artifacts in MRI. However, its effectiveness in improving the visibility of vessel-like thin structures in routine studies is unclear. In this study, we aim to demonstrate the ability of fat-navigator based PMC in improving the visibility of perivascular spaces (PVS) using data from two earlier studies.

**Methods:** Two open source MRI data set were used for motion artifact simulation and evaluating PMC, which consist of 66 T2-weighted images without PMC and 38 T2-weighted images with PMC. PMC was performed by adjusting field of view during scan based on motion parameters derived from fat navigators. Motion artifact simulation was performed by misplacing k-space data at a motion-related non-cartesian grid onto the cartesian grid calculated using motion-free images to generate the images without effects of PMC. The simulation’s ability to reproduce motion-induced blurring and ringing artifacts was evaluated using the sharpness at the lateral ventricle/white matter (WM) boundary and the magnitude of ringing artifact component in the Fourier spectrum. PVS volume fraction in WM was employed to reflect its visibility. Sharpness, magnitude of ringing artifact and PVS volume fraction were then compared between simulated images and real images with and without PMC.

**Results:** The consistencies in sharpness (rho ≥ 0.86, corrected p ≤ 4.4 ×10^-16^) and ringing artifact magnitude (rho ≥ 0.42, corrected p ≤ 0.001) were found between simulated images and real images without PMC. There was a significant negative correlation (rho ≤ -0.27, corrected p ≤ 0.08) between PVS volume fraction and motion severity in both simulated and real images without PMC. PMC removed the above correlations (rho ≥ -0.02, corrected p > 1) and increased the boundary sharpness compared to the images simulated using the same motion traces.

**Conclusions:** Motion artifact simulation can reproduce the desired motion-induced artifacts on images. PMC reduces the negative impacts of motion on image quality and improves PVS visibility.

## 1. Introduction

Perivascular spaces (PVS) are identified as the essential pathway for clearance of metabolic substance in glymphatic system [1], the abnormality of which has been found to be associated with a variety of neurovascular and neurodegenerative diseases [2]. Recent advances in high spatial resolution MRI allowed the morphological quantification of PVSs in vivo. However, as resolution improves, MR scans are prolonged and become more sensitive to head movements. Motion can induce blurring and ringing artifacts and noises in the reconstructed image [3, 4], rendering inaccuracy and unreliability of PVS measurement and reducing its diagnostic or scientific relevance [5].

To mitigate the motion-induced artifacts, two widely used schemes including retrospective motion correction (RMC) [5-9] and prospective motion correction (PMC) have been reported [10-18]. An earlier study [5] using fat navigators (FatNav) to estimate motion profile has found the beneficial effects of RMC on measuring PVS visibility and physiological changes. However, the effectiveness of FatNav -based PMC on improving PVS visibility remains unclear.

To evaluate the performance of PMC, uncorrected reference images are required to quantify image quality improvements. Several methods have been proposed to obtain the uncorrected images [10-17], which require additional or prolonged scans. Motion artifact simulation can avoid additional data acquisition and is therefore a more feasible approach for evaluating PMC in routine studies [18].

In this work, we proposed a motion simulation approach that takes into account acquisition process and uses motion-free multi-channel combined images and motion profiles derived from FatNav images. We studied the effects of motion and FatNav -based PMC on PVS visibility. First, we validated the veracity of motion artifacts in our simulation in terms of four parameters between simulated images and real images without PMC using different metrics for artifact level and PVS visibility. Then we studied the effects of real and simulated motion artifacts on WM and PVS visibility. Finally, we investigated the effects of PMC on image artifact level and PVS visibility by comparing real images with PMC with simulated images without PMC.

## 2. Materials and methods

### 2.1. Data acquisition and processing

#### 2.1.1 Participants

This study included two datasets from two earlier studies [19, 20]. The first dataset included 33 healthy volunteers (aged 21-55 years, 23 females). All subjects underwent a scan during air breathing and another scan during carbogen breathing, resulting in 66 images in total [19]. No motion correction was performed but navigators were acquired and will be named the NoPMC group in this study. The second dataset with the PMC-enabled, named PMC group in this study, included 19 participants with diabetes mellitus and 19 age- and sex-matched healthy controls (aged 34-70 years, 21 females) [20].

#### 2.1.2 MRI protocol

All images were acquired on a 7T MRI scanner (Siemens Healthineer, Erlangen, Germany) equipped with an 8-channel (NoPMC group) or a single channel (PMC group) transmitter and a 32-channel receiver head coil (Nova Medical, Wilmington, MA, USA). No B_1_ shimming for reducing radiofrequency (RF) field inhomogeneity was performed when using the 8-channel transmitter.

For both datasets, a 3D variable flip angle turbo spin echo (TSE) sequence was used to acquire T2 weighted images for imaging PVSs using the following parameters: TR/TE = 3000 (NoPMC) or 3300 (PMC)/326 ms, field of view (FOV) = 210 ×210 ×99.2 mm^3^, matrix size = 512 ×512 ×248, voxel size = 0.41 ×0.4 ×0.4 mm^3^, axial slices, TA = 8:03 (NoPMC) or 8:48 (PMC) min and all k-space data at a single partition encoding (PAR) step was acquired during each TR. Partial Fourier sampling was performed with a factor of 0.79 and 0.625 along the phase encoding (PE) and PAR directions, respectively. Undersampling factor was 3 with 24 auto-calibration lines along PE direction. Oversampling factor was 0.0323 along PAR direction.

A 3D FatNav [21] was embedded within each TR to monitor motion. With a binomial excitation pulse to selectively excite fat signal centered at 3.4 ppm upfield from water, the sequence parameters were set as follows: TR/TE = 3/1.31 ms, flip angle = 7°, undersampling factors = 4 ×4 and partial Fourier factor = 0.75 along both PE and PAR directions, FOV = 220 ×220 ×180 mm^3^ (NoPMC) or 222 ×198 ×210 mm^3^ (PMC), matrix size = 100 ×100 ×82 (NoPMC) or 74 ×66 ×70 (PMC), voxel size = 2.2 ×2.2 ×2.2 mm^3^ (NoPMC) or 3 ×3 ×3 mm^3^ (PMC), axial slices and TA = 0.89 s (NoPMC) or 0.47 s (PMC). A FatNav image with fully sampled rectangular region around k-space center was acquired before the first TR for obtaining calibration data for GRAPPA reconstruction.

#### 2.1.3 Prospective motion correction

Prospective motion correction works by dynamically modifying scanner parameters in real time according to motion data. During the scans, the FatNav images were reconstructed and then registered to the 1^st^ reference image using the vendor software installed on the scanner (MOCO functor) in order to yield relative motion parameters. These parameters were transmitted to the sequence to adjust the imaging FOV for the next TR such that the relative position between the imaged object and the FOV remained the same throughout the scan.

#### 2.1.4 Image reconstruction

The TSE images were reconstructed using vendor provided software on the scanner and the FatNav images were reconstructed using the GRAPPA algorithm [22].

### 2.2 Motion artifact simulation

Motion reduces image quality by introducing inconsistencies into k-space data. Due to motion, the MR signals corresponding to k-space data are placed into wrong positions in the k-space before inverse Fourier transformation (iFFT) into the image space. Therefore, the key of motion artifact simulation is to generate the k-space data at the real positions of readout lines in non-Cartesian grid, and then rearrange them into the intended Cartesian k-space coordinates.

The simulation included the following steps: (1) Due to oversampling along the PAR direction, we expanded the initial magnitude image by zero filling four voxels on each edge such that its Fourier transform matches the sampled k-space positions. (2) The three rotational motion parameters in each TR were converted into a 3×3 matrix *A* for transforming the intended Cartesian k-space coordinates *k* to the real measured non-Cartesian coordinates *k*′ by *k*′ = *Ak*. (3) The magnitude images expanded by (1) were then Fourier transformed into the k-space at the real k-space coordinates *k*′ through 3D non-uniform fast Fourier transform (NUFFT) [23]. (4) Then the k-space data were “incorrectly” placed onto the predefined Cartesian coordinates *k*. The phase at each k-space position was adjusted according to *S*′(*k*) = *S*(*k*′) · *e*^−*ik*·Δ*r*^ to account for the linear phase shift due to translational motion Δ*r*. The translational motion Δ*r* is related to the translational motion parameters Δ*r*_0_ derived from the FatNav registration by Δ*r* = −Δ*r*_0_ + (*A*^−1^ − *I*) · Δ*p*, where *I* is the unit matrix and Δ*p* is the difference between the TSE and FatNav imaging FOV centers. (5) Since the magnitude images we used possess only real components with zeros phases, the k-space data not acquired due to partial Fourier acceleration at coordinate *k* was filled by the complex conjugate of data at −*k*. (6) The motion-corrupted images were obtained by applying iFFT and then removing the four voxels at each edge of the FOV along PAR direction.

Our motion artifact simulation was performed using motion profiles of all scans in the NoPMC and PMC groups and two baseline images from subjects (named Subject 1 and Subject 2) in the NoPMC group with small motion scores (motion scores = 0.87 and 0.80 mm) and no apparent artifacts. The simulation was implemented in MATLAB 2021a (The MathWorks, Inc., Natick, Massachusetts, United States) and each image took approximately 11 min on a computer equipped with 3.3GHz Intel Xeon W-2275 CPU.

### 2.3 Data analysis

#### 2.3.1 Motion parameters

The six rigid-body motion parameters were obtained at each TR either using the 3dvolreg tool in AFNI [24] (NoPMC group) or the MOCO functor (PMC group), including rotation and translation around or along left-right (L-R), anterior-posterior (A-P) and superior-inferior (S-I) spatial directions. The motion profiles were estimated by registering the FatNav images onto the 1^st^ TR image and subtracted by motion parameters corresponding to time point at which the k-space center was acquired.

To quantify motion severity for each subject, motion score was calculated by combining the root sum square of translational and rotational motion ranges [5]. The motion severity was classified into four levels based on the motion score: no motion (motion score <= 0.9 mm), mild (0.9 mm < motion score <= 2 mm), moderate (2 mm < motion score <= 4mm) and severe (motion score > 4mm).

#### 2.3.2 Blurring artifact measurement: lateral ventricle/white matter boundary sharpness

Although the goal of the study is to study the effects of motion artifacts on PVS/WM contrast, as the image contrast between PVS and WM depends strongly on the partial volume effects which vary between PVS from different subjects in the simulated and real images [25], we instead choose to evaluate the effects of motion on image contrast using the sharpness [26] at the lateral ventricle/WM boundary which should have similar partial volume effects across subjects. Sharpness is quantified using the full width at half maximum (FWHM) of a Gaussian function whose integral is an s-shaped edge function. The edge function is used to fit the signal variation across tissue boundary. An increase in FWHM is indicative of increased blurriness and decreased image sharpness due to motion.

The region of interest (ROI) for edge function fitting was a cuboid across the lateral ventricular boundary drawn manually using ITK-SNAP version 3.8 [27] on the central sagittal slice with an average size of 7 (S-I) ×3 (L-R) ×6 (A-P) mm^3^, as shown in Fig. 1(a). The signal intensity versus the distances of all voxels to the boundary plane were then fitted by the edge function using the lsqcurvefit function in MATLAB 2020b (The MathWorks, Inc., Natick, Massachusetts, United States) to compute the FWHM.

**Figure 1.**
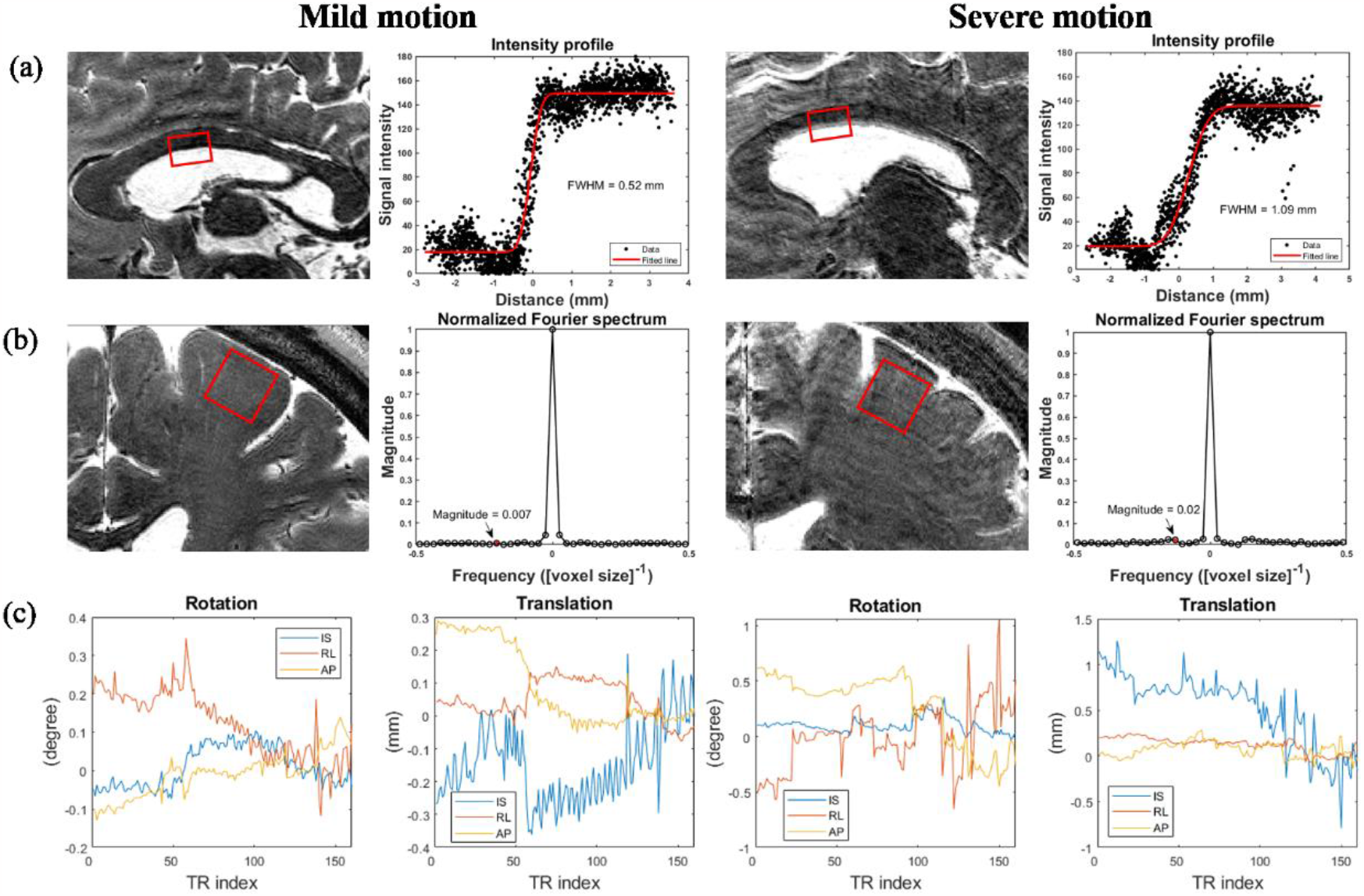
Examples of blurring and ringing artifact at mild and severe motion severities on cropped sagittal and coronal MIP slices of real NoPMC images. (a) are the lateral ventricle/white matter boundary sharpnesses calculated by the FWHMs of intensity profile estimated from red cuboid ROI. (b) are magnitudes of ringing artifacts in the Fourier spectrum. (c) are corresponding motion profiles, with motion scores of 1.25 and 4.19 mm, respectively.

#### 2.3.3 Ringing artifact measurement: magnitude of high-frequency component in Fourier spectrum

Ringing artifact generates oscillation around tissue boundaries, which manifests as high-frequency components in the frequency domain. Therefore, the ringing artifact can be quantified by the magnitude at the oscillation frequency. We used the closeness in magnitude at the oscillation frequency between the real and simulated images to evaluate the effectiveness of ringing artifact simulations.

We first determined the ROI in WM on a coronal slice with an average size of 16 (S-I) ×16 (L-R) mm^2^ where the stripes parallel to the brain surface were prominent as shown in Fig. 1(b), and averaged the signal intensity along the direction parallel to the brain surface. The resulting line intensity profile perpendicular to the stripes was then Fourier transformed to obtain its spectrum in the frequency domain. Then the magnitudes in spectrum were normalized to the range of 0 to 1. We identified the frequency of ringing artifact by counting the number of ringing cycles in the ROI, and then obtained the magnitudes at the peak position for both real and simulated images.

#### 2.3.4 Ghosting artifact measurement: background noise level

Ghosting artifact can be considered as a structured noise, which usually appears as a spatial shift of imaged object along the PE direction [28, 29]. The presence of ghosting artifact can be examined not only from the overlapping brain structure, but also from the background noise level. We determined the background noise level by the standard deviation of signal intensities in non-anatomical region. The non-anatomical region mask includes all voxels outside the skull which was segmented using FSL-FAST [30] from the magnitude images.

#### 2.3.5 PVS and WM visibility measurement

To quantitatively measure the PVS and WM visibility, a three-dimensional multi-channel multi-scale fully convolutional neural network (M^2^EDN) [31] was applied to delineate tubular PVSs and a classic encoder-decoder network (U-Net) [32] was applied to segment WM. The WM and PVS volume were calculated by multiplying the voxel size and the number of segmented voxels. The PVS volume fraction was calculated by the ratio of PVS volume to WM volume.

To clearly visualize vessel-like thin PVSs, the maximum intensity projection (MIP) with a 6 mm-thick section along the viewing A-P and S-I directions centered at the viewing slice was performed to project the hyperintense PVS signals onto the coronal and axial slices.

#### 2.3.6 Statistical analysis

FWHMs and ringing artifact magnitudes were not normally distributed for the NoPMC group in both real and simulated cases (p ≤ 0.007, Shapiro-Wilk tests). Real noise level was not normally distributed for the NoPMC group (p = 0.05, Shapiro-Wilk tests). Simulated FWHMs, real and simulated magnitudes were not normally distributed for the PMC group (p ≤ 1.58 ×10^-4^, Shapiro-Wilk tests). Due to non-normally distributed data, Spearman’s correlation test was performed to investigate the similarity between real and simulated motion artifacts.

Motion scores, WM volume, PVS volume and PVS volume fractions were not normally distributed for the NoPMC and PMC groups in both real and simulated cases (p ≤ 0.006, Shapiro-Wilk tests). To study the dependency of PVS visibility on *real* motion artifacts, Spearman’s partial correlation test was performed to study the relationship between PVS volume fraction and motion score, taking age (NoPMC) or age and disease (PMC) as confounding factors, both of which have been previously identified as factors that influence PVS visibility [25, 33]. To study the dependency of PVS visibility on *simulated* motion artifacts, Spearman’s correlation test between PVS volume fraction and motion score was performed.

The significance threshold was set to 0.05 for p-values after corrected by Bonferroni correction to account for multiple comparisons. The statistical analysis was performed using RStudio version 4.2.1 (RStudio, Inc., Boston, MA, USA).

## 3. Results

### 3.1 Performance of motion artifact simulation

In the NoPMC group, the motion scores ranged from 0.36 mm to 5.14 mm, with an average of 1.41 mm (± 1.03 mm). Of the 66 NoPMC images, the motion score distribution was: 29 (43.9%) belong to no motion, 22 (33.3%) belong to mild motion, 11 (16.7%) belong to moderate motion, and 4 (6.1%) belong to severe motion.

Two examples of mild and severe motion severities in the NoPMC group are shown in Fig. 1, with the motion scores of 1.25 and 4.19 mm calculated from motion profiles in Fig. 1(c), respectively. As the motion severity increased, FWHMs (0.52 and 1.09 mm) also increase as calculated from the image intensity profiles in Fig. 1(a), indicating the decreased sharpness at lateral ventricular boundary. Fig. 1(b) shows the magnitudes (0.007 and 0.02) of ringing artifacts in the normalized Fourier spectrum, suggesting the increased contribution of ringing artifacts to image quality deterioration at large motion.

Fig. 2 shows that the comparison results of blurring, ringing, and ghosting artifacts between simulated and real images. Figs. 2(a-c) and Figs. 2(d-f) are the results for Subjects 1 and 2, respectively. Figs. 2(a) and 2(d) show that the sharpnesses calculated on the real and the simulated images were roughly proportional to each other, with a proportionality constant from a linear fit with zero intercept close to identity (0.97 and 1.02, respectively, for Subjects 1 and 2). The Spearman’s correlation coefficients were 0.86 and 0.94 (corrected p ≤ 4.4 ×10^-16^), respectively. The weaker but still significant correlations were found in the spectral magnitude of ringing artifacts between the real and the simulated images, with the Spearman’s correlation coefficients of 0.48 and 0.42 (corrected p ≤ 0.001), respectively, as shown in Figs. 2(b) and 2(e). However, there were negative but non-significant correlations between the simulated and real background noise level as shown in Figs. 2(c) and 2(f). The negative and non-significant correlation was also found between the real noise level and motion score (rho = -0.03, p = 0.84, data not shown).

**Figure 2.**
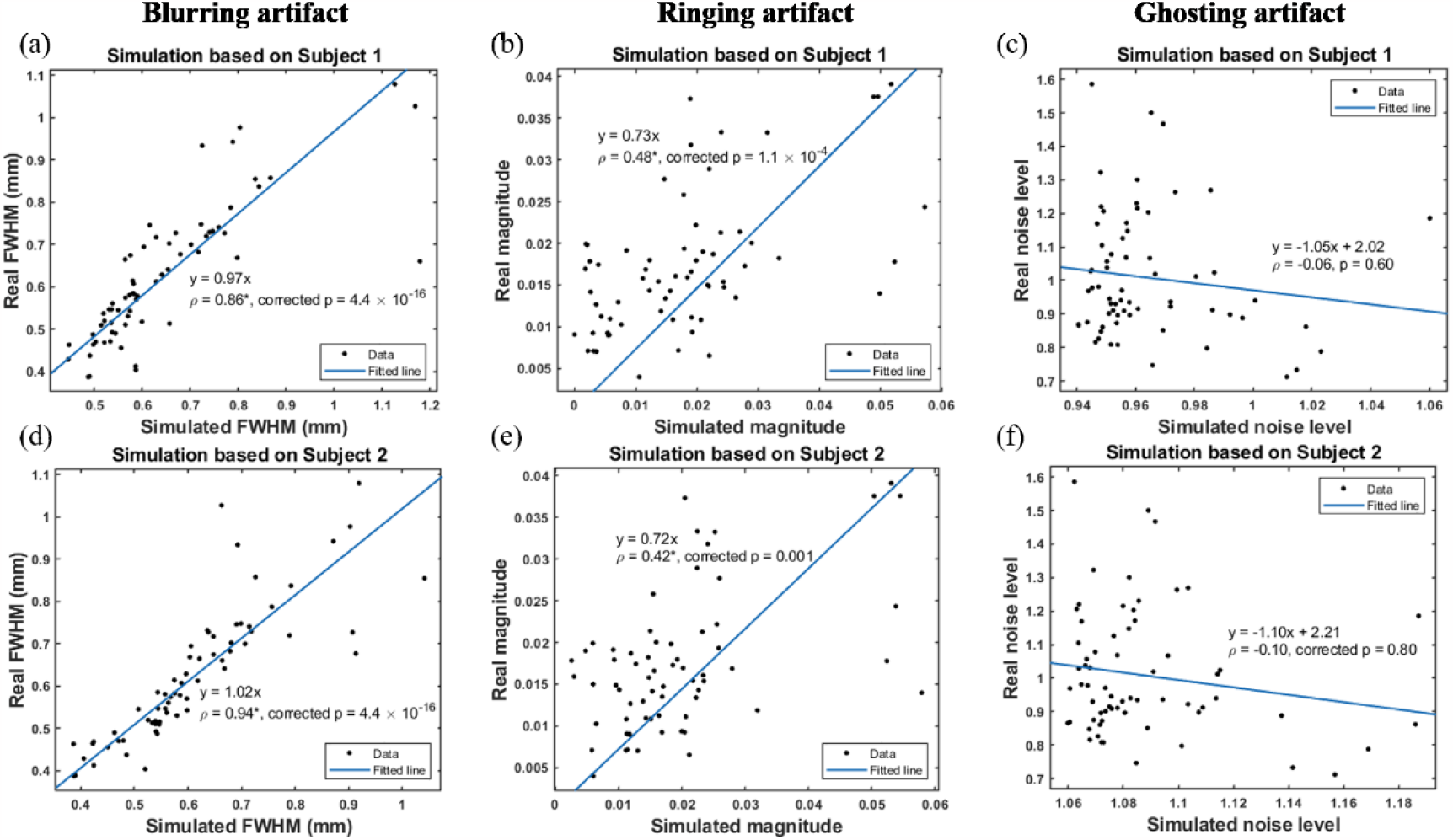
Comparison results in blurring, ringing and ghosting artifacts between the real images and the simulated images in the NoPMC group. Motion artifact simulations were conducted based on Subjects 1 (a-c) and 2 (d-f) and motion profiles of all real NoPMC images, respectively. Linear regression coefficients and Spearman correlation coefficients are shown, where asterisks next to the coefficients denote significance.

### 3.2 Effects of motion artifacts on WM and PVS visibility

To visualize the effects of *real* motion artifacts on PVS visibility at different motion severities with and without PMC, Fig. 3 and 4 show representative measured PVS images after MIP at different motion severities in the NoPMC and PMC groups, respectively. For the NoPMC group shown in Figs. 3(a-c), the motion scores of the three scans were 1.24, 2.38 and 5.10 mm, respectively, and the PVSs/WM boundary became blurrier as the motion score increased. Their motion traces are shown in Figs. 3(d-f). For the PMC group shown in Fig. 4, the motion scores of the three scans were 1.35, 2.27 and 4.95 mm, respectively, and the PVSs/WM boundary still remained clearly defined despite the presence of some ringing artifacts.

**Figure 3.**
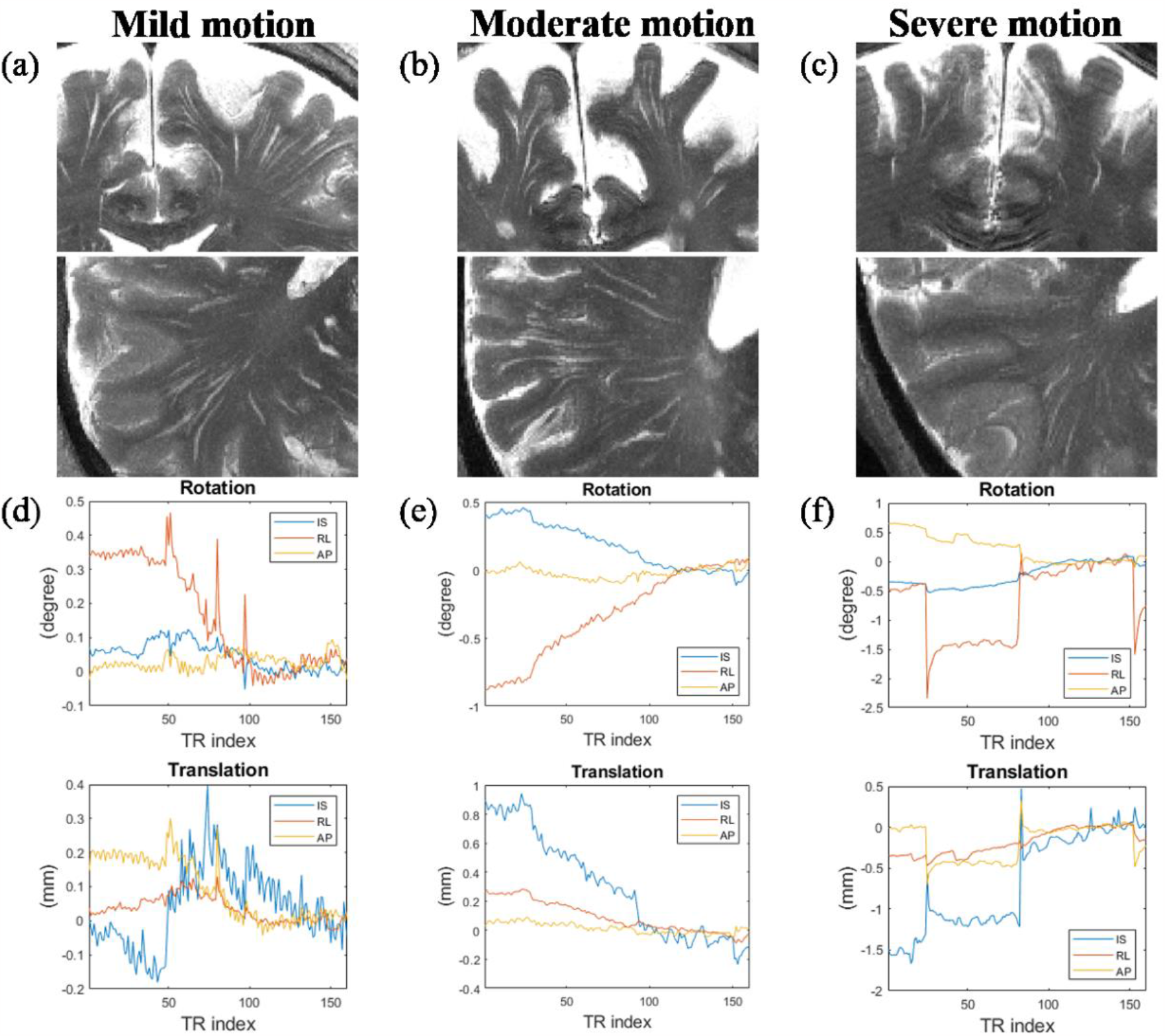
Examples of PVS visibility at different motion severities on cropped coronal and axial MIP slices of real NoPMC images. (a-c) are real NoPMC images, with motion scores of 1.24, 2.38 and 5.10 mm and PVS volume fractions of 1.32%, 0.59% and 0.56%, respectively. (d-f) are corresponding motion profiles.

**Figure 4.**
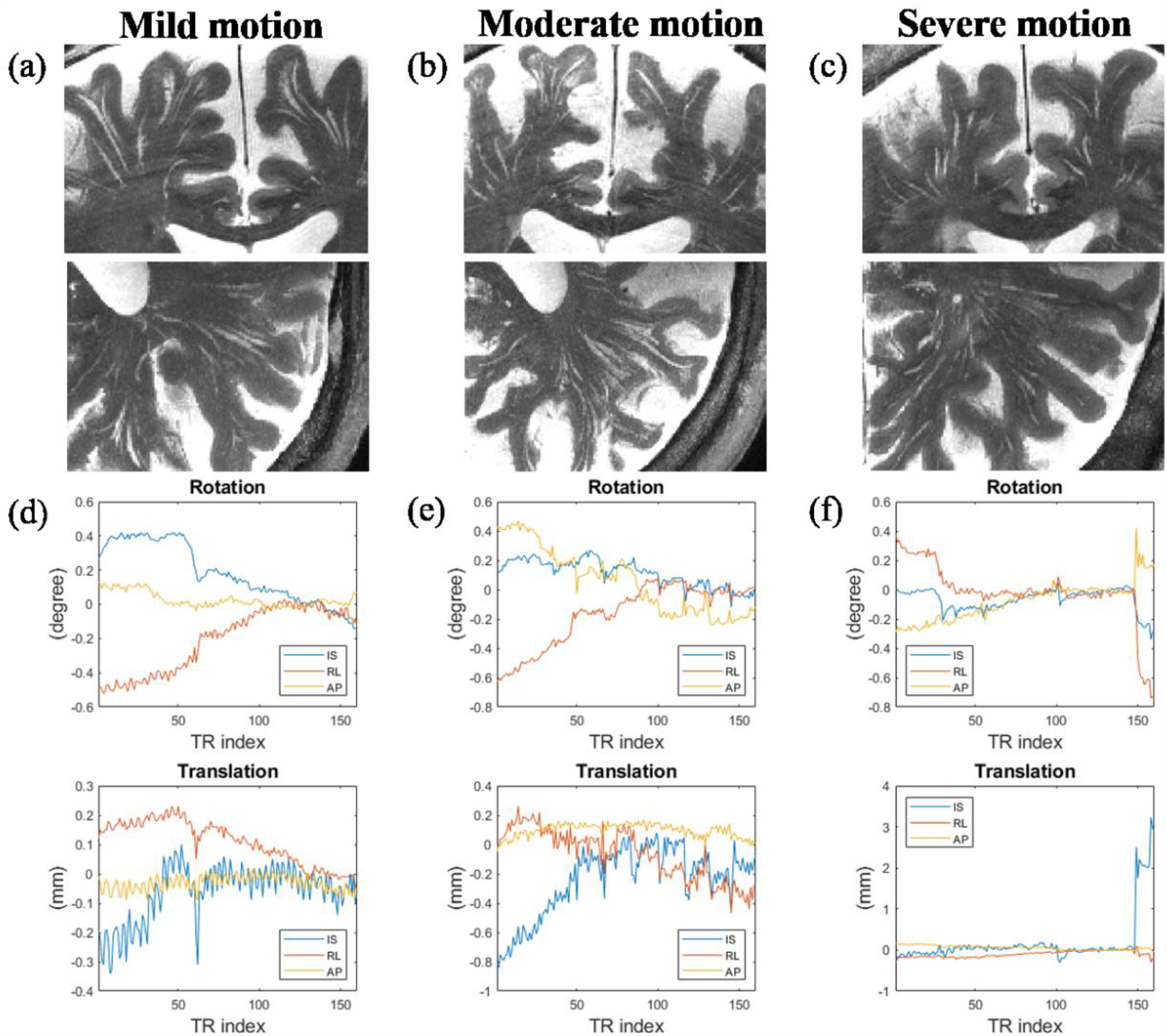
Examples of PVS visibility at different motion severities on cropped coronal and axial MIP slices of real PMC images. (a-c) are real PMC images, with motion scores of 1.35, 2.27 and 4.95 mm and PVS volume fractions of 1.40%, 2.06% and 1.20%, respectively. (d-f) are corresponding motion profiles.

To visualize the effects of *simulated* motion artifacts on PVS visibility at different motion severities, Figs. 5(b-d) and 5(f-h) display the MIP of the simulated images generated using the motion traces in Figs. 3(d-f) and in Figs. 4(d-f), and the baseline images of Subjects 1 and 2 (shown in Fig. 5a and 5e, respectively). There was a clear decrease of PVS visibility as the motion score increased in both groups as denoted by red arrows in Fig. 5, since the PMC effects was not considered in the simulation. The original volume fraction of Subject 1 (Subject 2) was 0.55% (1.05%). After simulation, the volume fractions became 0.46% (0.97%), 0.40% (0.85%) and 0.26% (0.82%), respectively, as the motion severity increased.

**Figure 5.**
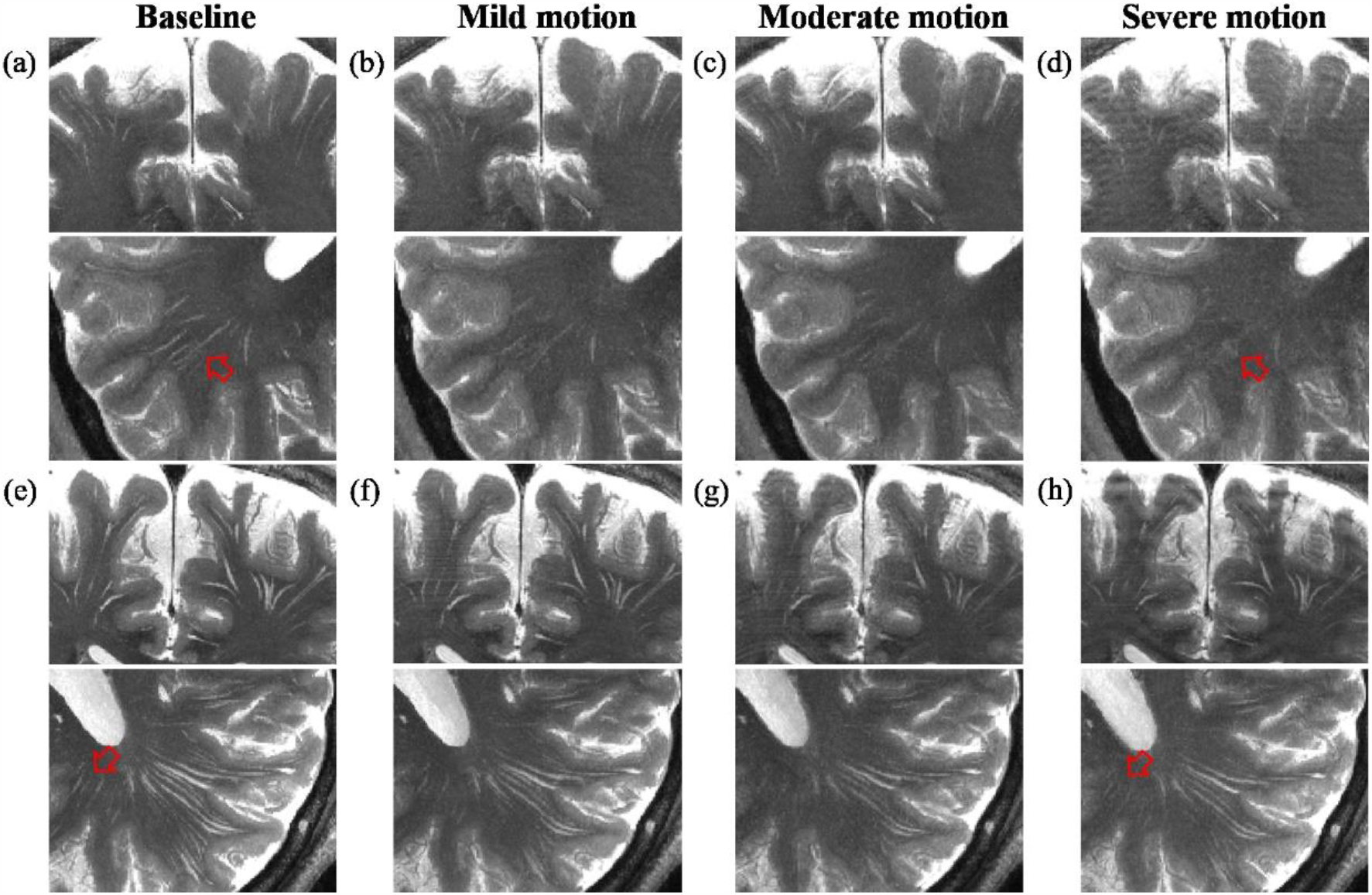
Examples of PVS visibility at different motion severities on cropped coronal and axial MIP slices of simulated images generated from the NoPMC and PMC groups. Simulations (b-d) were conducted based on Subject 1 as shown in (a) and the motion profiles from the NoPMC images in Fig. 3(d-f), with motion scores of 1.24, 2.38 and 5.10 mm, respectively. Simulations (f-h) were conducted based on Subject 2 as shown in (e) and motion profiles from the PMC images in Fig. 4(d-f), with motion scores of 1.35, 2.27 and 4.95 mm, respectively.

Figs. 6(a-c) shows the scatter plots of the WM volume versus motion score on the real and simulated images, respectively. The negative correlations for the real NoPMC (blue points in Fig. 6a; rho = -0.03) and simulated images based on Subject 2 (Fig. 6c; rho = -0.53) were found, while no significant correlation was found for the simulated images based on Subject 1 (Fig. 6b; rho = 0.14).

**Figure 6.**
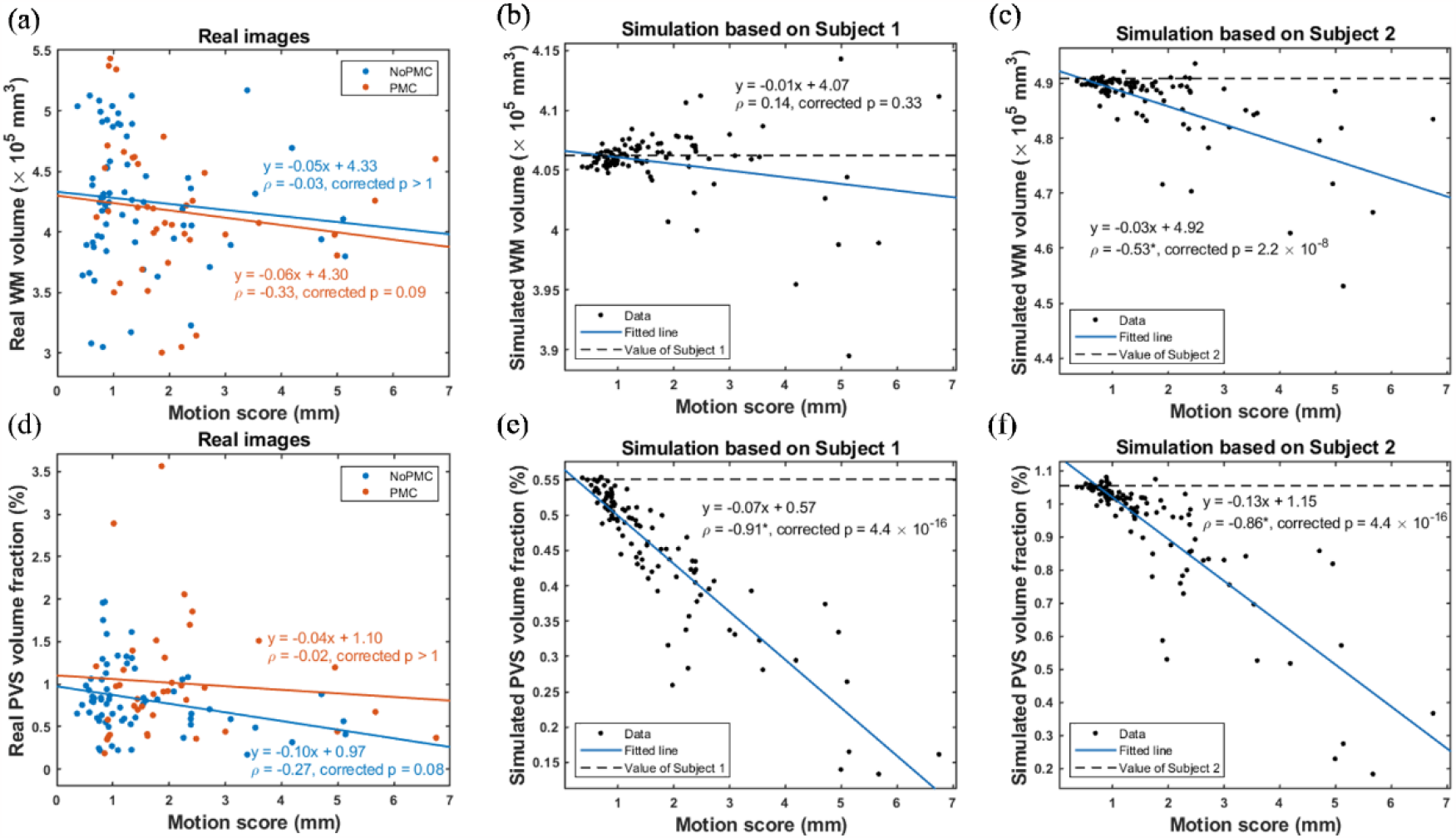
Effects of real and simulated motion artifacts on WM and PVS visibility. (a-c) shows the scatter plot of WM volume versus motion score. (d-f) shows the scatter plot of PVS volume fraction versus motion score. The dotted lines in b-c and e-f refer to the values calculated from the baseline image (Subjects 1 and 2). Linear regression coefficients and Spearman correlation coefficients are shown (Spearman partial correlation coefficients are shown in d), where asterisks next to the coefficients denote significance.

Figs. 6(d-f) shows the scatter plots of the PVS volume fraction versus motion score on the real and simulated images, respectively. The negative correlations between PVS volume fraction and motion score were significant in the simulated images (Figs. 6e and 6f; rho ≤ -0.86, corrected p ≤ 4.4 ×10^-16^), and marginally significant in the real NoPMC images (blue points in Fig. 6d; rho = -0.27, corrected p = 0.08) prior to Bonferroni correction. In contrast to the WM volume, the correlations of PVS volume fraction were much stronger in both real NoPMC and simulated images, but the correlation was weaker and non-significant on the real PMC images (rho ≥ -0.02, corrected p > 1).

### 3.3 Performance of prospective motion correction

In the PMC group, the motion scores ranged from 0.70 mm to 6.75 mm, with an average of 2.12 mm (± 1.37 mm). Of the 38 PMC images, the motion score distribution was: 3 (7.9%) belong to no motion, 21 (55.3%) belong to mild motion, 10 (26.3%) belong to moderate motion, and 4 (10.5%) belong to severe motion.

Fig. 7 compares the blurring artifacts, ringing artifacts and PVS visibility between NoPMC and PMC groups. The real PMC images had lower FWHMs (0.51 ± 0.07 mm) than the real NoPMC images (0.62 ± 0.15 mm), as shown in Figs. 7(a) and 7(c). In contrast to the NoPMC group, there was no significant correlation in the FWHMs between the real PMC images and the simulated images (p ≥ 0.37) and the proportionality constants from a linear fit with zero intercept were less than 1 (0.57 and 0.65, respectively, for Subjects 1 and 2). The FWHM of the real PMC image was on average reduced by 0.35 and 0.45 mm compared to the simulated images based on Subjects 1 and 2, respectively, suggesting improved image sharpness after PMC. Figs. 7(b) and 7(e) compare the magnitudes of ringing artifacts obtained from real versus simulated images in the NoPMC and PMC groups. There were weaker correlations (rho ≤ 0.19, corrected p ≥ 0.48) in the PMC group compared to the NoPMC group (rho ≥ 0.42, corrected p ≤ 0.001). Figs. 7(c) and 7(f) compare the PVS volume fractions calculated from real versus simulated images in the NoPMC and PMC groups. After accounting for factors (age and disease condition) that affect visibility, significant correlations in volume fractions between real and simulated images were observed only in the NoPMC group (corrected p ≤ 0.009), but not in the PMC group (p ≥ 0.79).

**Figure 7.**
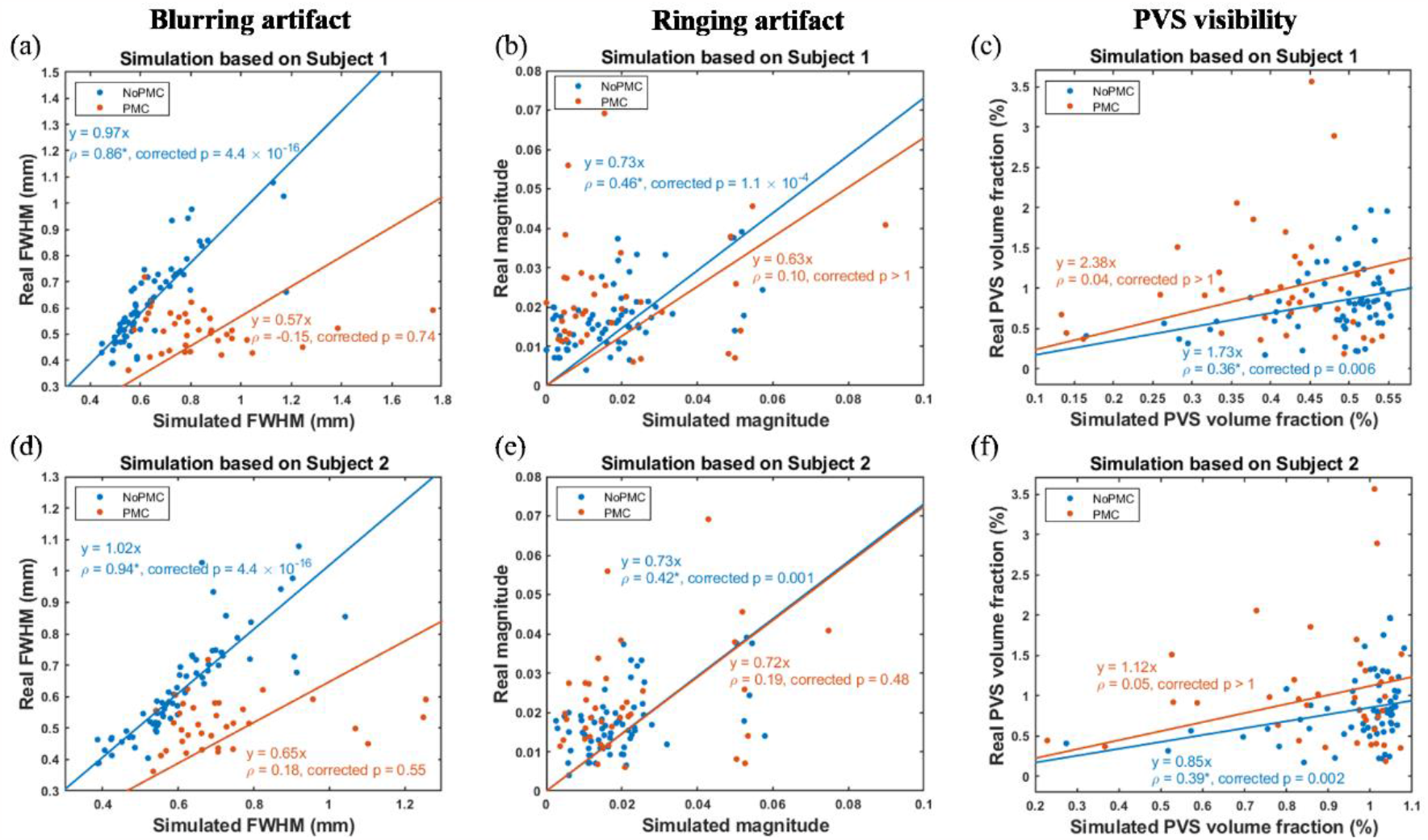
Comparison results in blurring artifacts, ringing artifacts and PVS visibility on the real images and the simulated images between the NoPMC and PMC groups. Motion artifact simulations were conducted based on Subjects 1 (a-c) and 2 (d-f) and motion traces of all real NoPMC and PMC images, respectively. Linear regression coefficients and Spearman correlation coefficients are shown, where asterisks next to the coefficients denote significance.

## 4. Discussion

In this study, we proposed motion artifact simulation as a mean to measure the effects of PMC on PVSs. This method allows for generation of uncorrected images to evaluate the PMC performance without the need of MR scan repetition. We validated the effectiveness of simulation in unintentional real motion scenarios and evaluated its ability in simulating blurring and ringing artifact. We found that the simulated motion artifacts degraded the PVS visibility as the real artifacts did. We found that PMC can improve image sharpness and mitigate such adverse effects on PVS visibility.

Several methods have been proposed to obtain the uncorrected images in order to assess the performance of PMC, including multiple scans with PMC-enabled and -disabled [10-16], single scan with interleaved acquisition of PMC-enabled and - disabled [17], and motion artifact simulation [18]. However, to evaluate the effectiveness of PMC in routine clinical studies, only motion artifact simulation is feasible. Multiple scans and interleaved acquisition will prolong the scan time that affects the comfort level of subjects and increases the likelihood of head movements. Furthermore, in methods using multiple scans, the motion observed in each scan cannot be identical though usually the subjects are instructed to perform deliberate movements [13-16]. In addition, the effectiveness of PMC on correcting intentional motion-induced artifacts may not be generalizable to routine studies where more complex unintentional motion can occur.

Multiple methods have been proposed for performing motion artifact simulation [34], including those based on magnitude images [6-9] and those based on multi-channel complex k-space data [18, 35]. Motion effects were accounted for by either modifying k-space coordinates while keeping the data intact [8, 18, 35], or by modifying the k-space data at the same coordinates[6, 7, 9].

Magnitude image-based methods require less computational time and is applicable to more studies as the raw k-space data are not always available. Besides, modifying k-space data as adopted in our study matches more closely the actual data sampling and image reconstruction process in the presence of motion. In such approach, the simulated images are expected to contain artifacts only resulting from k-space coordinate misplacements and phase inconsistencies due to the measured motion. In contrast, in addition to such artifacts, the reverse RMC images also contain artifacts due to other motion-related effects in the actual data, such as B0 homogeneity changes and uncaptured fast motion.

The PVS volume fraction in real NoPMC images showed a non-significant negative correlation with motion score (Fig. 6d) but a stronger significant correlation with the simulation PVS volume fraction (Figs. 7c and 7f), suggesting that our motion simulation can more accurately reflect the effects of motion on the PVS visibility than the motion score as the artifact level depends on not only motion range but also motion pattern. Furthermore, we found the significant correlations in the blurring and ringing artifact between simulated and real NoPMC images, further demonstrating the veracity of the simulated motion-induced artifacts. All the correlations were reduced (Fig. 7) if the real NoPMC images were replaced by the PMC images, thus suggesting the effectiveness of PMC in mitigating the negative effect of motion on PVS visibility.

In contrast to the PVS volume fraction, the correlation of WM volume with motion score were weaker in the simulated cases (Figs. 6b-c), which suggests lower sensitivity of WM visibility to motion than the smaller thin-tubular structures like PVS. In addition, we found the non-significant correlations between simulated and real background noise level (Figs. 2c and 2f), suggesting the dominance of other noise sources such as thermal and physiological noises.

Several studies have identified PVS as a potential biomarker of neurovascular and neurodegenerative diseases [2, 36-39]. PVS play a crucial role in the removal of metabolic substances and waste products from brain. Its abnormalities can lead to the accumulation of toxins and wastes, which may contribute to the pathogenesis of these diseases. The improvements in PVS visibility and robustness against motion on MRI by PMC can facilitate the investigation of PVS abnormalities, especially in longitudinal studies, which may provide novel insights into the underlying neuropathogenesis and promote the development of therapeutic targets.

There are some limitations of the presented motion artifact simulation. First, no real images with and without motion from the same subjects were used for validating the simulation method. Second, although the computation demand can be considerably reduced, the simulation bias conducted on the k-space of magnitude image or the raw k-space data remain to be studied. Furthermore, the simulation neglected the potential impact of magnetic field inhomogeneity and RF transmitting and receiving coil sensitivity, which might change due to head position and orientation during scan. The resulting MR signal variations could lead to deviations of simulated artifacts.

## 5. Conclusions

In conclusion, our proposed motion artifact simulation provides an effective tool for investigating the effects of motion and PMC on PVS visibility. In the presence of motion, the FatNav-based PMC was shown to mitigate the negative impacts of motion and increase the number of MRI-visible PVSs in WM. Our study demonstrated the value of PMC for improving the quality of PVS images in investigations aimed at further illuminating the potential role of PVS abnormalities in the pathogenesis of neurodegenerative disorder.

